# Cell Cycle, Energy Metabolism and DNA Repair Pathways in Cancer Cells are Suppressed by Compound Kushen Injection

**DOI:** 10.1101/348102

**Authors:** Jian Cui, Zhipeng Qu, Yuka Harata-Lee, Thazin Nwe Aung, Hanyuan Shen, Wei Wang, David L Adelson

## Abstract

In this report we examine candidate pathways perturbed by Compound Kushen Injection (CKI) a Traditional Chinese Medicine (TCM) that we have previously shown to alter the gene expression patterns of multiple pathways and induce apoptosis in cancer cells. We have measured protein levels in HEPG2 and MDA-MB-231 cells for genes in the cell cycle pathway, DNA repair pathway and DNA double strand breaks (DSBs) previously shown to have altered expression by CKI. We have also examined energy metabolism by measuring [ADT]/[ATP] ratio (cell energy charge), lactate production and glucose consumption. Our results demonstrate that CKI can suppress protein levels for cell cycle regulatory proteins and DNA repair while increasing the level of DSBs. We also show that energy metabolism is reduced based on reduced glucose consumption and reduced cellular energy charge. Our results validate these pathways as important targets for CKI. We also examined the effect of the major alkaloid component of CKI, oxymatrine and determined that it had no effect on DSBs, a small effect on the cell cycle and increased the cell energy charge. Our results indicate that CKI likely acts through the effect of multiple compounds on multiple targets where the observed phenotype is the integration of these effects and synergistic interactions.

## Introduction

Compound Kushen Injection (CKI) is a complex mixture of plant bioactives extracted from Kushen (*Sophora flavescens*) and Baituling (*Smilax Glabra*) that has been approved for use in China since 1995 by the State Food and Drug Administration (SFDA) of China (State medical license no. Z14021231). CKI is widely used in China as an adjunct for both radiotherapy and chemotherapy in cancer. While most of the data supporting its use have been anecdotal and there is little clinical trial data demonstrating its efficacy, it has been shown to be effective at reducing sarcoma growth and cancer pain in an animal model[1] and cancer pain in patients [2].

CKI contains over 200 chemical compounds including alkaloids and flavonoids such as matrine, oxymatrine and kurarinol, and has previously been shown to affect the cell cycle and induce apoptosis in cancer cells[1, 2, 3; 4, 5, 6, 7]. Furthermore, functional genomic characterisation of the effect of CKI on cancer cells using transcriptome data indicated that multiple pathways were most
likely affected by CKI [4]. These observations support a model wherein many/all of the individual compounds present in CKI can act on many single targets or on multiple targets to induce apoptosis.

Based on previously reported work [4] and our currently unpublished work (Cui *et al*)[8], specific pathways were selected for follow up experiments to validate their response to CKI in order to formulate more specific hypotheses regarding the mechanism of action of CKI on cancer cells. We had previously shown that CKI altered the cell cycle and induced apoptosis while altering the expression of many cell cycle genes in three cancer cell lines [4, 8]. We had also shown that DNA repair pathway genes were significantly down-regulated by CKI and that energy production related to NAD(P)H synthesis from glycolysis and oxidative phosphorylation was reduced by CKI. As a result we focused on the following candidate pathways: cell cycle, DNA repair and glucose metabolism to validate their alteration by CKI. We used two cell lines for these validation experiments, one relatively insensitive to CKI (MDA-MB-231) and one sensitive to CKI (HEPG2). Furthermore, while the literature shows varying effects for major compounds present in CKI on cancer cells [9, 10], we also tested oxymatrine, the major alkaloid found in CKI and widely believed to be very important for the effects of CKI, on our selected pathways.

## Materials and methods

### Cell culture and chemicals

CKI with a total alkaloid concentration of 26.5 mg/ml in 5 ml ampoules was provided by Zhendong Pharmaceutical Co. Ltd. (Beijing, China). Cell culture methods have been previously described [4].

A human breast adenocarcinoma cell line, MDA-MB-231 and a hepatocellular carcinoma cell line HEPG2 were purchased from American Type Culture Collection (ATCC, Manassas, VA). The cells were cultured in Dulbecco’s Modified Eagle Medium (DMEM; Thermo Fisher Scientific, MA, USA) supplemented with 10% foetal bovine serum (Thermo Fisher Scientific). Both cell lines were cultured at 37°C with 5% CO2. For all *in vitro* assays, cells were cultured overnight before being treated with either CKI (at 1 mg/ml and 2 mg/ml of total alkaloids). As a negative control, cells were treated with medium only and labelled as “untreated”. After 24 and 48 hours of treatment, cells were harvested and subjected to the downstream experiments.

All the *in vitro* assays employed either 6-well plates or 96-well plates. The seeding density for 6-well plates for both cell lines was 4×10^5^ cells and treatment methods were as previously described [4]. The seeding density of HEPG2 cells for 96 well plates was 4×10^3^ cells per well and for MDA-MB-231 cells was 8×10^4^ cells per well, and used the same treatment method as above: after seeding and culturing overnight, cells were treated with 2 mg/ml CKI diluted with complete medium for the specified time.

### Glucose consumption assay

Glucose consumption was assessed in both cell lines in 6-well plates. Glucose consumption was determined by using a glucose oxidase test kit (GAGO-20, Sigma, St. Louis, MO). After culturing for different durations (3, 6, 12, 24 and 48 hours), 50 *μ*l of culture medium was collected from untreated groups and treated groups. The cells were tripsinized for cell number determination using trypan blue exclusion assay and the number of bright, viable cells were counted using a hemocytometer. Collected suspension, blank medium and 2 mg/ml CKI, were all filtered and diluted 100 fold with MilliQ water. The absorbance at 560 nm was converted to glucose concentration using a 5*μ*g/ml glucose standard from the kit as a single standard. Glucose consumption was calculated by subtracting the blank medium value from treated/collected medium values. Glucose consumption per cell was calculated from the number of cells determined above.

### Measurement of [ADP]/[ATP] ratio

Cells were cultured in white 96-well plates with clear bottoms. The [ADP]/[ATP] ratio of both cell lines was determined immediately after the incubation period (24 and 48 hours) using an assay kit (MAK135; Sigma Aldrich, USA) according to the manufacturer’s instructions. Levels of luminescence from the luciferase-mediated reaction was measured using a plate luminometer (PerkinElmer 2030 multilabel reader, USA for CKI experiments or Promega, USA for oxymatrine experiments). The [ADP]/[ATP] ratio was calculated from the luminescence values using a formula provided by the kit manufacturer.

### Lactate content assay

The concentration of lactate, the end product of glycolysis, was determined using a lactate colorimetric assay kit (Abcam, Cambridge, MA, USA). Cells were cultured in 6-well plates, and then harvested and deproteinized according to the manufacturer’s protocol. The optical density was measured at 450 nm and a standard curve plot (nmol/well vs. OD 450 nm) was generated using serial dilutions of lactate. Lactate concentrations were calculated with formula provided by the kit manufacturer.

### Cell cycle assay

Cells were cultured in 6-well plates and treated with 2 mg/ml CKI or 0.5 mg/ml oxymatrine. After culturing for 3, 6, 12, 24 and 48 hours, cells were harvested and subjected to cell cycle analysis by Propidium Iodide staining as previously reported [4]. Data were obtained by flow cytometry using Accrui (BD Biosciences, NJ, US) and analysed using FlowJo software (Tree Star Inc, Ashland, Oregon, USA).

### Microscopy

After culturing for 48 hours on 8 well chamber slides, control and treated cells were fixed in 1% paraformaldehyde for 10 minutes at room temperature, washed with Phosphate Buffered Saline three times and permeabilized with 0.5% Triton X-100 for 10 minutes. After fixation and permeabilization, cells were blocked with 5% Fetal Bovine Serum for 30 minutes. Permeabilized cells were stained with 5*μ*g/ml of Alexa Fluor®594 conjugated anti-H2AX.X Phospho (red) (Biolegend, Ser139) in 5% Fetal Bovine Serum overnight followed with Alexa Fluor®488 Phalloidin (green) (Biolegend) staining for 20 minutes at 4°C.

Stained cells were mounted with 4’,6-diamidino-2-phenylindole (DAPI) and visualized with an Olympus FV3000 (Olympus Corporation, Tokyo, Japan) confocal microscope using a 60× oil objective. Fluorescence intensity was quantified using Imaris software (Bitplane, Saint Paul, MN) and averaged using at least 10 cells in each experiment.

### Flow-cytometry quantification of protein expression

Cells were cultured in 6-well plates and treated with CKI. After 24 and 48 hours, cells were harvested to detect intranuclear/intracellular levels of proteins involved in cell cycle and DNA DSBs pathways using the following antibodies; (cell cycle primary antibodies: (Cell Signaling Technologies, Danvers, MA, USA: P53 Rabbit mono-Ab, CCND1 Rabbit mono-Ab, CDK2 Rabbit mono-Ab) (Abcam, Cambridge, UK:CDK1 Rabbit mono-Ab), for these primaries, cell cycle isotype control: Cell Signaling Technologies: Rabbit igG, cell cycle secondary antibody Anti-rabbit (PE conjugated), additional primary antibody: CTNNB1 Rabbit mono-Ab (Alexa Fluor®647 conjugated) and isotype control: Rabbit IgG (Alexa Fluor®647 conjugated), both from Abcam)(DSBs antibody: Anti-H2AX (PE conjugated) primary antibody and isotype control: Mouse IgG1 (PE conjugated) both from BioLegend, San Diego, CA, USA) (DNA repair antibodies: primary antibodies - KU70 Rabbit mono-Ab (Alexa Fluor®647 conjugated) and KU80 Rabbit mono-Ab (Alexa Fluor®647 conjugated), Isotype control: Rabbit IgG (Alexa Fluor®647 conjugated), all from Abcam) Cells were sorted and the data acquired on a FACS Canto (BD Biosciences, NJ, US) or Accrui (BD Biosciences, NJ, US), and the data were analysed using FlowJo (Tree Star Inc.) software.

### Cell cycle functional enrichment re-analysis

In order to identify the phases of the cell cycle affected by CKI, differentially expressed gene data from [4] was submitted to the Reactome database[11], and used to identify functionally enriched genes.

### Statistical analysis

All measurements above were performed in triplicate and repeated at least three times. Statistical significance was determined by two-way ANOVA test; error bars represent standard deviation.

## Results

### Pathway validation

Based on our previous results indicating that CKI could suppress NAD(P)H synthesis [4] and (Supplementary Material, Figure S1), we examined the effect of CKI on energy metabolism by measuring glucose uptake, [ADP]/[ATP] ratio and lactate production. We measured glucose uptake in both CKI treated and untreated cells from 0 to 48 hrs after treatment and observed a reduction in glucose uptake (Fig. 1A). The growth curves for both cell lines were relatively flat after CKI treatment, in contrast to untreated cells. MDA-MB-231 cells, which are less sensitive to CKI in terms of apoptosis, had a higher level of glucose uptake than HEPG2 cells, which are more sensitive to CKI. Because the overall glucose uptake was consistent with the cell growth curves, the glucose consumption per million cells for each cell line and treatment was different. Untreated HePG2 cells maintained a relatively flat rate of glucose consumption per million cells, while for CKI treated HEPG2 cells the rate of glucose consumption per million cells decreased with time, becoming significantly less towards 48 hrs. The glucose consumption variance for both untreated and treated MDA-MB-231 cells was high, but both overall glucose consumption and glucose consumption per million cells appeared to decrease over time.

**Figure 1.**
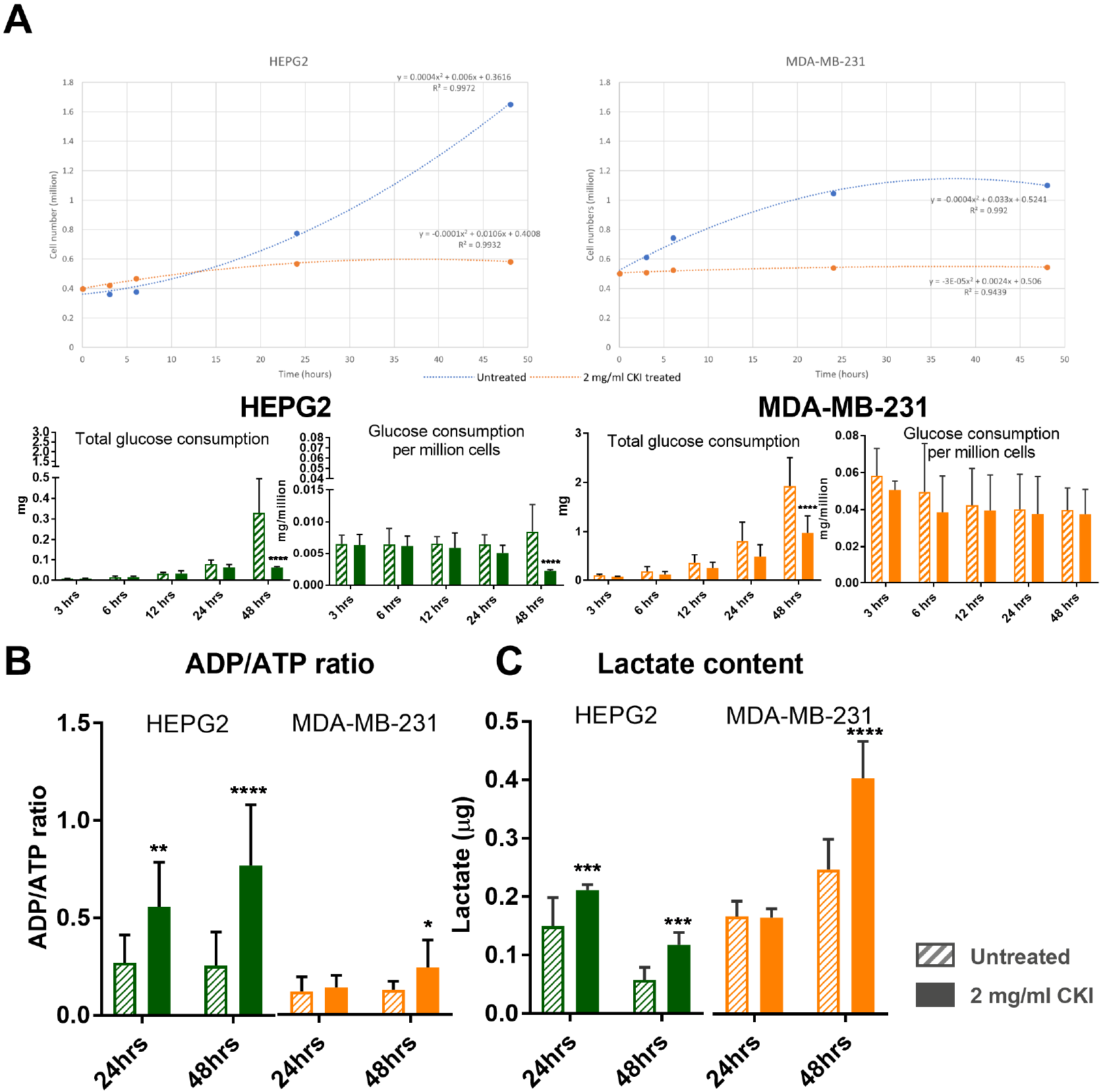
The energy metabolism determination assays in the two cell lines. A. Comparison of glucose consumption analysis between the two cell lines at 3, 6, 12, 24 and 48 hours. Overall glucose consumption is divided by cell number to calculate the consumption of glucose per million cells. B. [ATP]/[ADP] ratio assay result for the two cell lines at 24 and 48 hours. C. Lactate content detection for the two cell lines at 24 and 48 hours. Statistical analyses were performed using two-way ANOVA comparing treated with untreated (*p<0.05, **p<0.01, ***p<0.001, ****p<0.0001); bars show 1 standard deviation from the mean.

Because changes in glucose consumption are mirrored by other aspects of energy metabolism, we assessed the energy charge of both CKI treated and untreated cells by measuring the [ADP]/[ATP] ratio at 24 and 48 hours after treatment (Fig. 1B). HEPG2 cells had a lower energy charge (higher [ADP]/[ATP] ratio) compared to MDA-MB-231 cells and after CKI treatment both cell lines showed a decrease in energy charge, consistent with our previous measurements using a 2,3-Bis(2-methoxy-4-nitro-5-sulfonyl)-2H-tetrazolium-5-carboxanilideinner salt (XTT) assay (Supplementary Material, Figure S1). However the decrease in energy charge was earlier and much more pronounced for HEPG2 cells compared to MDA-MB-231 cells.

The flip side of glucose consumption is the production of lactate via glycolysis, which is the initial pathway for glucose metabolism. We therefore measured lactate production in order to determine if the observed decreases in energy charge and glucose consumption were directly attributable to reduced glycolytic activity. We measured intracellular lactate concentration in both CKI treated and
untreated cells at 24 and 48 hours after treatment (Fig. 1C) and found that lactate concentrations increased as a function of CKI treatment in both cell lines. This result is consistent with a build up of lactate due to an inhibition of the Tricarboxylic Acid (TCA) cycle leading to decreased oxidative phosphorylation and lower cellular energy charge. CKI must therefore inhibit cellular energy metabolism downstream of glycolysis, most likely at the level of the TCA cycle. Decreased energy charge can have widespread effects on a number of energy hungry cellular processes involved in the cell cycle, such as DNA replication.

Having validated the effect of CKI on cellular energy metabolism, we proceeded to examine the perturbation of cell cycle and expression of cell cycle proteins, as these are energy intensive processes. We had previously identified the cell cycle as a target for CKI based on transcriptome data from CKI treated cells [4, 8]. We carried out cell cycle profiling on CKI treated and untreated cells using Propidium Iodide staining and FACS (Fig. 2A) as described in Materials and Methods. The two cell lines displayed slightly different profiles to each other, but their response to CKI was similar in terms of an increase in the proportion of cells in G1-phase. For HEPG2 cells, CKI caused consistent reductions in the proportion of cells in S-phase accompanied by corresponding increases in the proportion of cells in G1-phase. This is indicative of a block in S-phase leading to accumulation of cells in G1-phase. For MDA-MB-231 cells, CKI did not promote a significant decrease in the proportion of cells in S-phase, but did cause an increase in the percentage of cells in G1 phase at 24hrs and a pronounced decrease in cells in G2/M phase at 12 hours.

**Figure 2.**
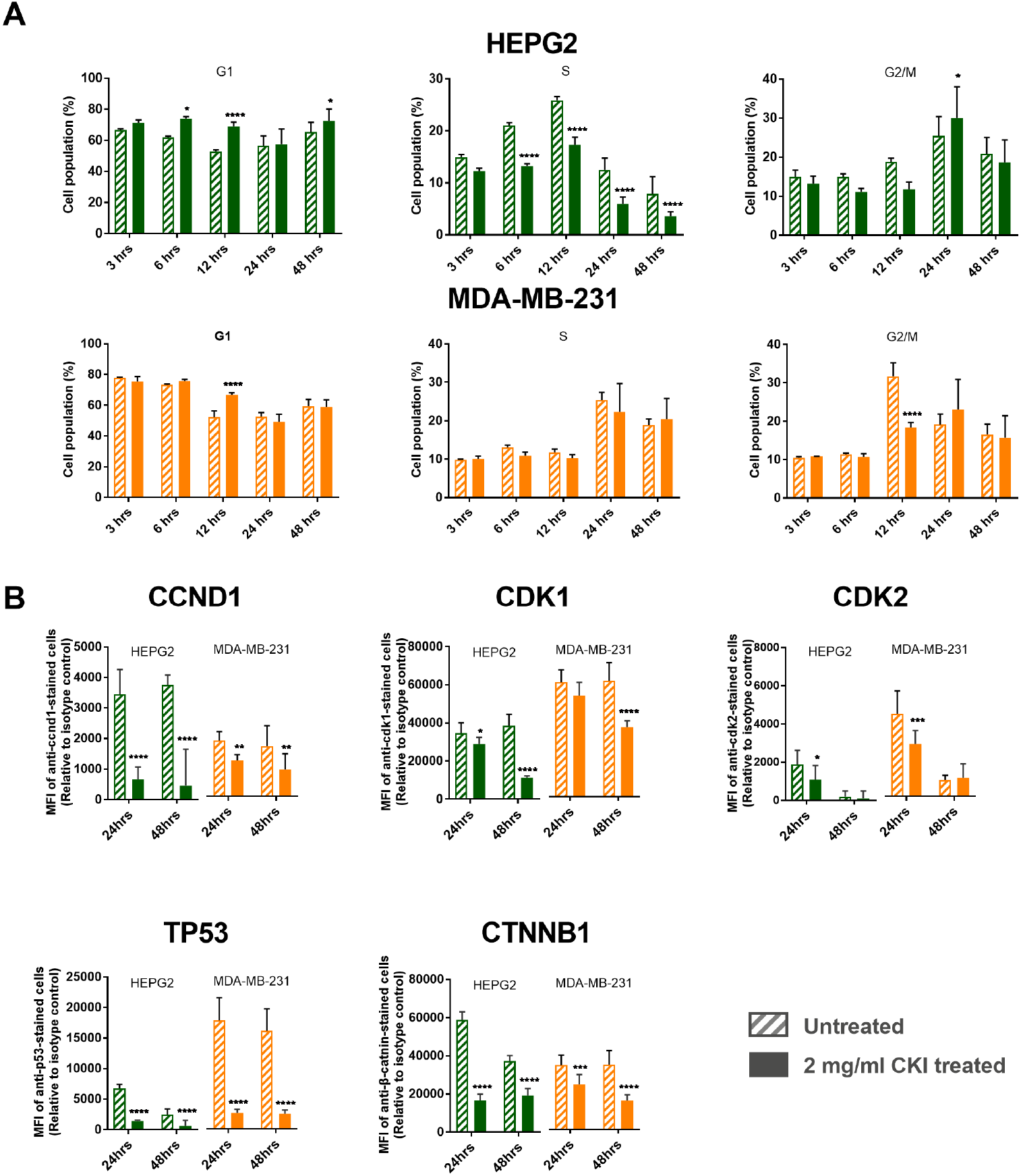
Cell cycle shift by CKI and undulating expression of the key proteins. A. Cell cycle shift regulated by CKI over 48 hours. In both cell lines, the earliest shifted cell cycle phase was S phase 6 hours after treatment. Compared to HEPG2, MDA-MB-231 showed delayed responses. B. Expression levels for 5 proteins (ccnd1, cdk1, cdk2, p53 and catenin *β* 1) as a result of CKI treatment at both 24 hours and 48 hours. Statistical analyses were performed using two-way ANOVA comparing with untreated (*p<0.05, **p<0.01, ***p<0.001, ****p<0.0001); bars show 1 standard deviation from the mean.

We also examined the levels of key proteins involved in the cell cycle pathway (Cyclin D1:CCND1,Cyclin Dependent Kinase 1:CDK1, Cyclin Dependent Kinase 2:CDK2, Tumor Protein p53:TP53 and Catenin Beta 1:CTNNB1) at 24 and 48 hours after CKI treatment previously shown to have altered transcript expression by CKI (Fig. 2B). Both cell lines showed similar results for all five proteins, with decreased levels caused by CKI, and validated previous RNAseq data [4, 8]. CCND1 regulaes the cell-cycle during G1/S transition. CDK-1 promotes G2-M transition, and regulates G1 progress and G1-S transition. CDK-2 acts at the G1-S transition to promote the E2F transcriptional program and the initiation of DNA synthesis, and modulates G2 progression. TP53 acts to negatively regulate cell division. CTNNB1 acts as a negative regulator of centrosome cohesion. Down-regulation of these proteins is therefore consistent with cell cycle arrest/disregulation and the cell cycle result in Fig. 2A. These results indicate that CKI alters cell cycle regulation consistent with cell cycle arrest. Cell cycle arrest is also an outcome that can result from DNA damage such as DNA double strand breaks (DSBs) [12].

We had previously observed that DNA repair genes had lower transcript levels in CKI treated cells [4, 8], so hypothesised that this might result in increased numbers of DSBs. We measured the expression of γ-H2AX in both cell lines (Fig. 3A) and found that it was only over-expressed at 48 hours in CKI treated HEPG2 cells. We also carried out localization of γ-H2AX using quantitative immunofluorescence microscopy and determined that the level of γ-H2AX increased in nuclei of CKI treated cells in both cell lines (Fig. 3B). These results indicated an increase in DSBs as a result of CKI treatment. In order to confirm if reduced expression of DNA repair proteins was correlated with the increase in DSBs we measured levels of Ku70/ku80 proteins in CKI treated cells (Fig. 3C). In HEPG2 cells, Ku80, a critical component of the Non-Homologous End Joining (NHEJ) DNA repair pathway was significantly down regulated at both 24 and 48 hours after CKI treatment. In MDA-MB-231 cells, Ku70 expression was down-regulated at both 24 and 48 hours after CKI treatment, and Ku80 was down-regulated at 24 hours. Because Ku70/Ku80 are subunits of a required DNA repair complex, reduced expression of either subunit will result in decreased DNA repair. Our results therefore support a suppressive effect of CKI on DNA repair, likely resulting in an increased level of DSBs.

**Figure 3.**
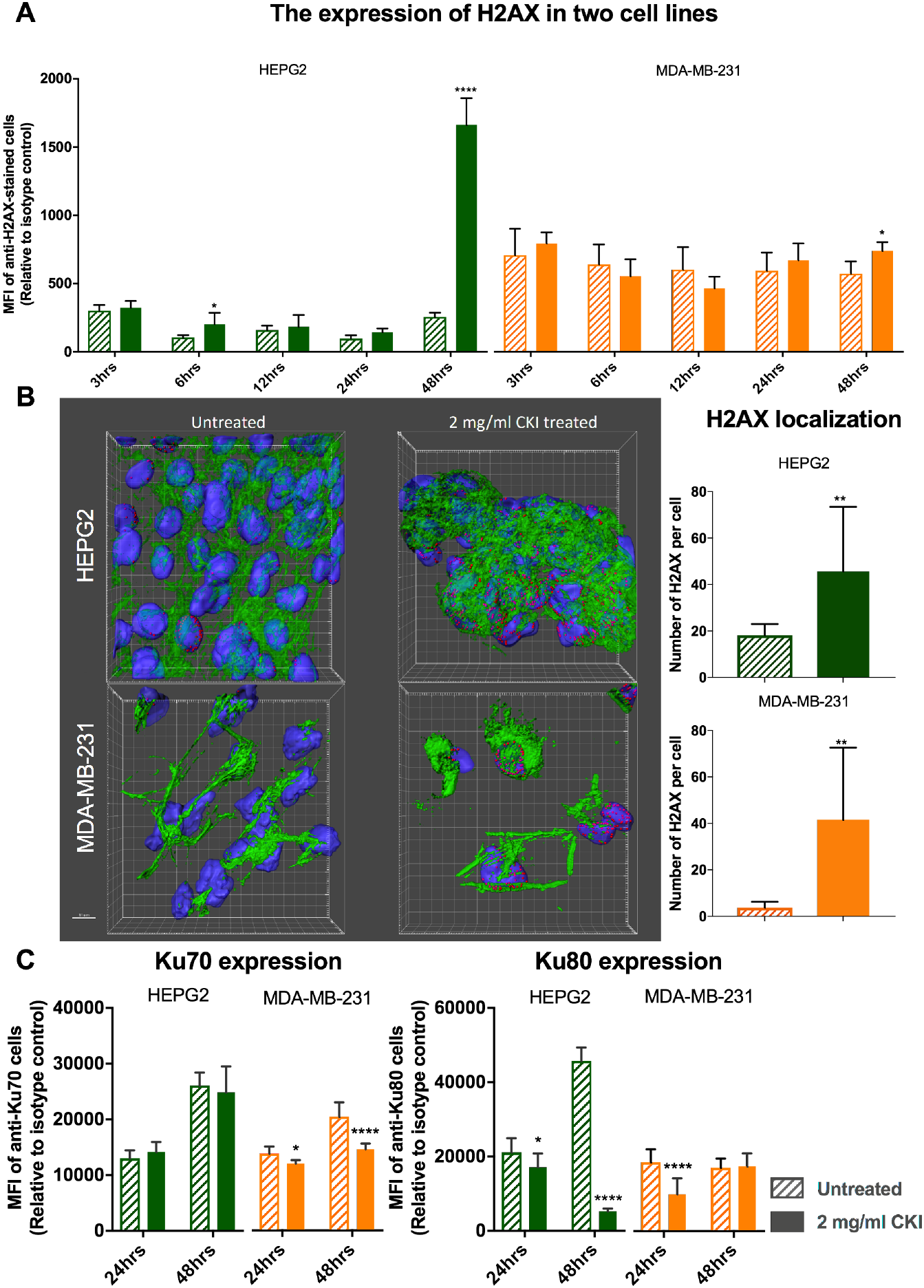
DSBs were increased by CKI treatment. A. γ-H2AX expression from 3 hours to 48 hours after treatment with 2 mg/ml CKI in two cell lines. B. Localization of γ-H2AX in two cell lines after CKI treatment for 48 hours. Green shows the cytoskeleton stained with an antibody to F-actin, blue is DAPI staining of nuclei, pink/red is staining of DSBs with antibody to γ-H2AX. The bar graph shows a quantification of the average number of γ-H2AX foci per cell detected in immunofluorescence images of 2 mg/ml CKI treated and untreated groups. At least 10 images of 3 independent replicate experiments were analyzed. C.Expression of DSBs repair proteins, Ku70 and Ku80, as a result of treatment with 2 mg/ml CKI in two cell lines. Statistical analyses were performed using two-way ANOVA comparing treated with untreated (*p<0.05, **p<0.01, ***p<0.001, ****p<0.0001); bars show 1 standard deviation from the mean.

### Effect of Oxymatrine, the principal alkaloid in CKI

Because CKI is a complex mixture of many plant secondary metabolites that may have many targets and there is little known about its molecular mode of action, we examined the effects of the most abundant single compound found in CKI, oxymatrine, on the most sensitive cell line, HEPG2. Oxymatrine is an alkaloid that has previously been reported to have effects similar to CKI, so we expected it might have an effect on one or more of our three validated pathways.

Oxymatrine, at 0.5 mg/ml which is equivalent to the concentration of oxymatrine in 2mg/ml CKI,
did not have an equivalent effect on the cell cycle compared to CKI (Fig. 4A vs Fig. 2A). Oxymatrine caused only minor changes to the cell cycle with small but significant increases in the proportion of cells in G1-phase at 3 and 48 hours and a small but significant decrease in the proportion of cells in S1-phase at 48 hours. Oxymatrine also caused a significant increase in the proportion of cells undergoing apoptosis in HEPG2 cells, albeit at a lower level than CKI (Supplementary Material, Figure 2).

**Figure 4.**
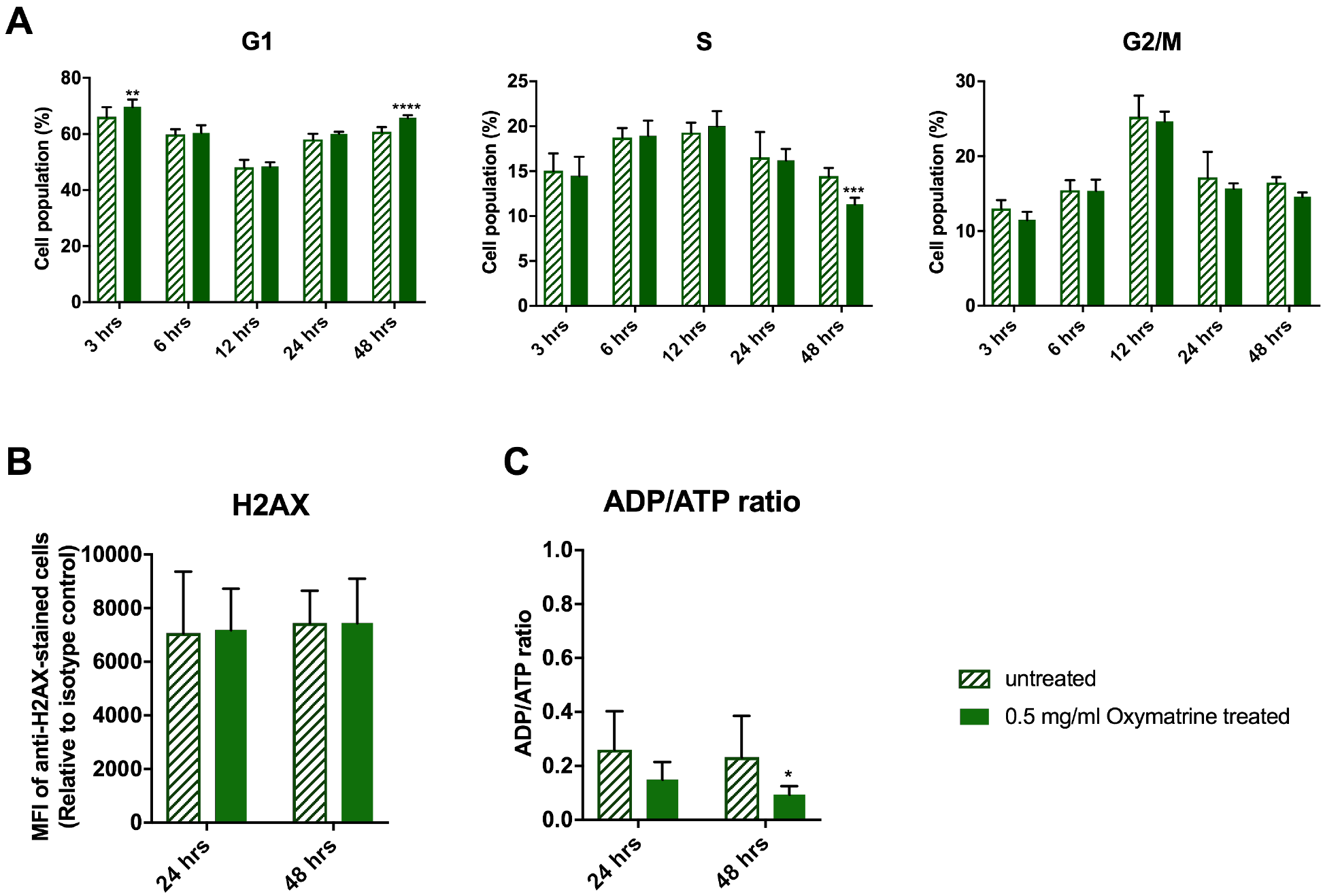
Effect of oxymatrine alone on validated pathways. Oxymatrine was tested at 0.5mg/mL which is equivalent to its concentration in CKI. A. Effect of oxymatrine on cell cycle in HEPG2 cells over 48 hours. B. Effect of oxymatrine on γ-H2AX (DSBs) levels after 24 and 48 hours. C. Effect of oxymatrine on [ADP]/[ATP] ratio after 24 and 48 hours. Statistical analyses were performed using two-way ANOVA comparing treated with untreated (*p<0.05, **p<0.01, ***p<0.001, ****p<0.0001); bars show 1 standard deviation from the mean.

Oxymatrine had no effect on H2AX levels in HEPG2 cells (Fig. 4B). This was in stark contrast to the effect of CKI (Fig. 3A) at 48 hours and indicated that oxymatrine alone had no effect on the level of DSBs.

Surprisingly, oxymatrine had the opposite effect on energy metabolism compared to CKI, causing a decrease in [ADP]/[ATP] ratio indicating a large increase in the energy charge of the cells (Fig. 4C).

### Integration of results

The effect of CKI on cancer cells was validated in all three of our candidate pathways: cell cycle, energy metabolism and DNA repair. Because these pathways are not isolated, but instead are integrated aspects of cell physiology CKI may act through targets in some or all of these three pathways or may act through other targets that either directly or indirectly suppress these pathways. CKI may also act through the synergistic effects of multiple compounds on multiple targets in our candidate pathways. This possibility is consistent with the partial and minor effects of oxymatrine on our candidate pathways.

## Discussion

We have validated three pathways (cell cycle, energy metabolism and DNA repair) that are perturbed by CKI and that can be used as the focus for further investigations to identify specific molecular targets that mediate the perturbations.

### Cell cycle perturbation by CKI

Our results show that CKI can perturb the cell cycle by altering the proportions of cells in G1-phase, S-phase and G2/M-phase. This result is similar to what we have observed before [4, 8], but has not been widely reported in the literature. The alkaloid oxymatrine, the most abundant compound present in CKI, has also been shown previously to perturb a number of signaling pathways [13] and alter/arrest the cell cycle in a variety of cancer cells [14, 15, 16, 17, 18] and we have confirmed this observation (Fig. 4A) in HEPG2 cells. Our results permit direct comparison with CKI because our experiments have been done using eqivalent concentrations of oxymatrine alone or in CKI. While oxymatrine has an effect on the cell cycle, it is not as effective at perturbing the cell cycle as is CKI. This indicates that oxymatrine must interact with other compounds in CKI to have a stronger effect on the cell cycle.

### Energy metabolism suppression by CKI

We have shown for the first time that CKI can inhibit energy metabolism as demonstrated by lower levels of NADH/NADPH and a higher [ADP]/[ATP] ratio. These results, combined with lower glucose utilisation and higher lactate levels indicate that this suppression was likely due to inhibition of the TCA cycle or oxidative phosphorylation. Previously, Gao *et al* [3] have reported that CKI significantly increased the concentration of pyruvate in the medium and this observation in combination with our results supports a decrease in metabolic flux through the TCA cycle as the likely cause of the reported suppression of energy metabolism. Interestingly, oxymatrine on its own had the opposite effect on [ADP]/[ATP] ratio compared to CKI, indicating that it can enhance energy metabolism and increase the energy charge of the cell.

### DNA repair suppression by CKI

There is only one report in the literature of oxymatrine inducing DSBs [19] and no reports with respect to CKI. Our results show for the first time that not only does CKI induce DSBs, but that is also likely inhibits DNA repair by decreasing the expression of the Ku70/Ku80 complex required for NHEJ mediated DNA repair. It is worth noting however, that the reported effect of oxymatrine on DSBs [19] uses significantly higher (4-8 fold) concentrations of oxymatrine compared to our experiments. In our hands oxymatrine alone at 0.5mg/ml showed no effect on DSBs as judged by the level of γ-H2AX after 24 or 48 hours.

## Conclusions

CKI causes suppression of energy metabolism and DNA repair along with altered cell cycle (summarized in Fig. 5). CKI has also previously been reported to induce apoptosis in cancer cells [4]. The overarching question is if CKI has independent effects on these three pathways or if the primary effect of CKI is through a single pathway that propagates effects to other, physiologically linked pathways. It may be that CKI suppresses energy metabolism thus disrupting downstream, energy hungry processes such as DNA replication and DNA repair. Alternatively, there could be independent effects on DNA repair leading to checkpoint induced cell cycle perturbation/arrest. Our results based on oxymatrine treatment of HEPG2 cells indicate that the cell-cycle is likely directly affected by oxymatrine and thus CKI. However oxymatrine alone had no effect on DNA repair and boosted, rather than reduced the energy charge of the cell. Taken together, these results support a model of many compounds/many targets [20] for the mode of action of CKI, where multiple compounds affect multiple targets and the synergistic, observed effect is significantly different to that seen with individual components.

**Figure 5.**
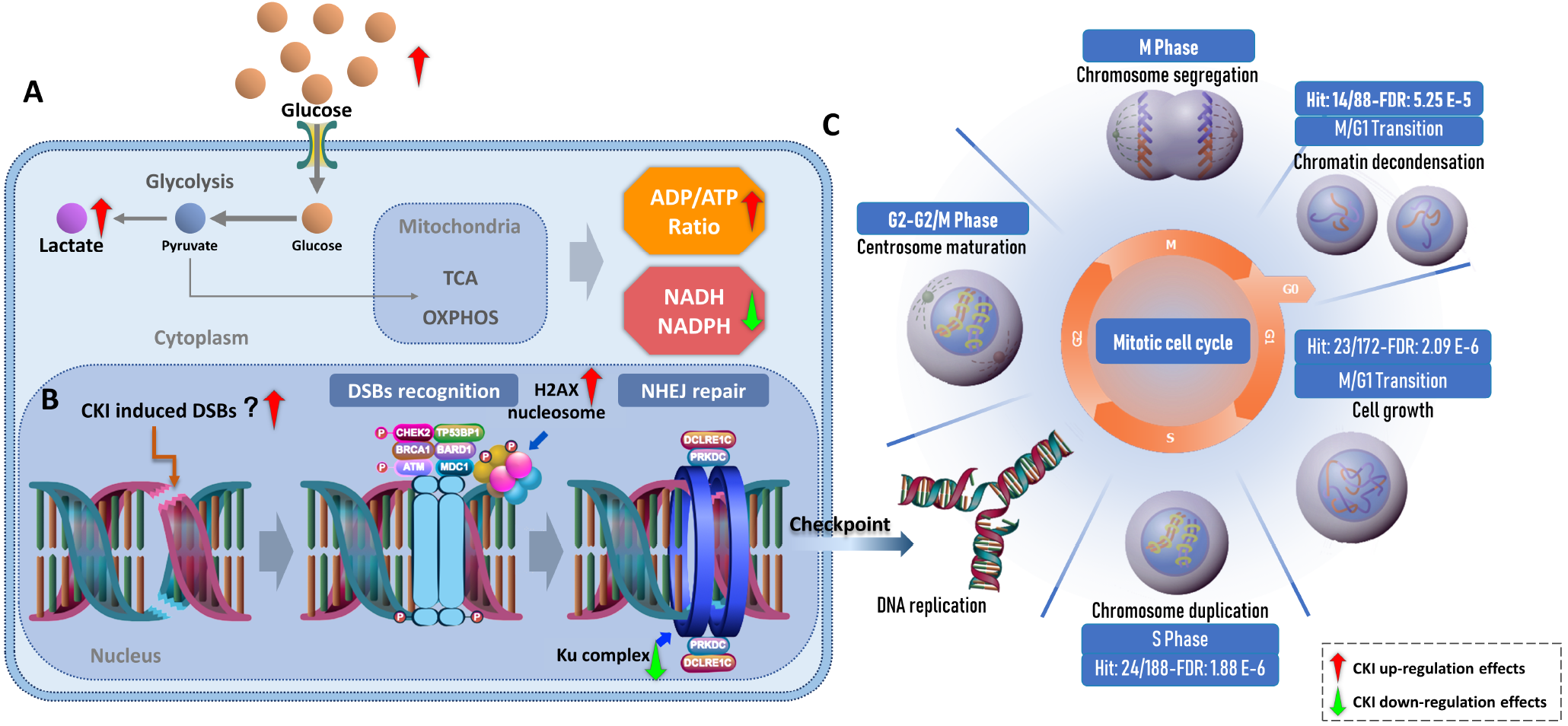
Integration of the three pathways altered by CKI. A. General presentation of energy metabolism affected by CKI. Glucose utilization is down-regulated by CKI. This is accompanied by increased lactate in the cytoplasm as CKI inhibits glucose metabolism downstream of glycolysis, leading to an increase in [ADP]/[ATP] and decrease in NADH/NAPDH. B. Effects on DNA repair in cancer cells by CKI. CKI may be able to directly induce DSBs, but CKI may also indirectly induce DSBs by arresting checkpoint functions during the cell cycle. In addition, CKI may also inhibit NHEJ, the major repair mechanism for DSBs. C. Reactome functional enrichment of cell cycle genes based on shared differentially expressed (DE) genes from previous studies. From M/G1 to S phase, the shared DE genes from both cell lines were significantly enriched. Most of these DE genes, were down-regulated.

## Acknowledgements

We would like to thank Prof. Frank Grutzner for providing DAPI, Adelaide Microscopy for training and equipment use and Jue Zhang, Bo Han and Dan Kortschak for helpful discussions.

## Availability of data and materials

All data analyzed in this study are available from the public source NCBI (https://www.ncbi.nlm.nih.gov/) and details can be found in the supplementary file.

## Author’s contributions

J.C, Z.Q., Y.H-L. and D.L.A. designed research; J.C., Y-H-L., Z.Q., T.N.A., H.S. and W.W. performed research; and J.C., Z.Q., Y.H-L. and D.L.A wrote the paper.

## Ethics approval and consent to participate

Not applicable.

## Consent for publication

Not applicable.

## Additional Files

### Additional file 1 — Supplementary Information

Additional file contains supplementary figures and tables as referred to in the main body of the paper.

## Abbreviations

DSBs: double strand breaks
Ku70/Ku80: the Ku heterodimer proteins

## References

1. Zhao Z, Fan H, Higgins T, Qi J, Haines D, Trivett A, et al. Fufang Kushen injection inhibits sarcoma growth and tumor-induced hyperalgesia via TRPV1 signaling pathways. Cancer Letters. 2014 Dec;355(2):232–241. Available from: http://dx.doi.org/10.1016/j.canlet.2014.08.037.

2. Wang W, You Rl, Qin Wj, Hai Ln, Fang Mj, Huang Gh, et al. Anti-tumor activities of active ingredients in Compound Kushen Injection. Acta Pharmacologica Sinica. 2015;36(6):676.

3. Gao L, Wang KX, Zhou YZ, Fang JS, Qin XM, Du GH. Uncovering the anticancer mechanism of Compound Kushen Injection against HCC by integrating quantitative analysis, network analysis and experimental validation. Sci Rep. 2018 Jan;8(1):624.

4. Qu Z, Cui J, Harata-Lee Y, Aung TN, Feng Q, Raison JM, et al. Identification of candidate anti-cancer molecular mechanisms of Compound Kushen Injection using functional genomics. Oncotarget. 2016 10;7(40):66003–66019.

5. Zhou SK, Zhang RL, Xu YF, Bi TN. Antioxidant and immunity activities of Fufang Kushen Injection Liquid. Molecules. 2012 May;17(6):6481–90.

6. Sun M, Cao H, Sun L, Dong S, Bian Y, Han J, et al. Antitumor activities of kushen: literature review. Evid Based Complement Alternat Med. 2012;2012:373219.

7. Xu W, Lin H, Zhang Y, Chen X, Hua B, Hou W, et al. Compound Kushen Injection suppresses human breast cancer stem-like cells by down-regulating the canonical Wnt/β-catenin pathway. J Exp Clin Cancer Res. 2011 Oct;30:103.

8. Cui J, Qu Z, Harata-Lee Y, Shen H, Aung TN, Wang W, et al. The Effect of Compound Kushen Injection on Cancer Cells: Integrated Identification of Candidate Molecular Mechanisms.;. (Manuscript submitted).

9. Ma X, Li RS, Wang J, Huang YQ, Li PY, Wang J, et al. The Therapeutic Efficacy and Safety of Compound Kushen Injection Combined with Transarterial Chemoembolization in Unresectable Hepatocellular Carcinoma: An Update Systematic Review and Meta-Analysis. Frontiers in Pharmacology. 2016 Mar;7. Available from: https://dx.doi.org/10.3389/fphar.2016.00070.

10. Zhongquan Z, Hehe L, Ying J. Effect of compound Kushen injection on T-cell subgroups and natural killer cells in patients with locally advanced non-small-cell lung cancer treated with concomitant radiochemotherapy. Journal of Traditional Chinese Medicine. 2016;36(1):14–18. Available from:http://www.sciencedirect.com/science/article/pii/S0254627216300024.

11. Fabregat A, Jupe S, Matthews L, Sidiropoulos K, Gillespie M, Garapati P, et al. The Reactome pathway knowledgebase. NUCLEIC ACIDS RESEARCH. 2018;46(D 1):D649–D655.

12. Shaltiel IA, Krenning L, Bruinsma W, Medema RH. The same, only different - DNA damage checkpoints and their reversal throughout the cell cycle. J Cell Sci. 2015 Feb;128(4):607–20.

13. Lu ML, Xiang XH, Xia SH. Potential Signaling Pathways Involved in the Clinical Application of Oxymatrine. Phytotherapy Research. 2016 May;30(7):1104–1112. Available from: http://dx.doi.org/10.1002/ptr.5632.

14. Li W, Yu X, Tan S, Liu W, Zhou L, Liu H. Oxymatrine inhibits non-small cell lung cancer via suppression of EGFR signaling pathway. Cancer Med. 2018 Jan;7(1):208–218.

15. He M, Jiang L, Li B, Wang G, Wang J, Fu Y. Oxymatrine suppresses the growth and invasion of MG63 cells by up-regulating PTEN and promoting its nuclear translocation. Oncotarget. 2017 Sep;8(39):65100–65110.

16. Li S, Zhang Y, Liu Q, Zhao Q, Xu L, Huang S, et al. Oxymatrine inhibits proliferation of human bladder cancer T24 cells by inducing apoptosis and cell cycle arrest. Oncol Lett. 2017 Jun;13(6):4453–4458.

17. Wu J, Cai Y, Li M, Zhang Y, Li H, Tan Z. Oxymatrine Promotes S-Phase Arrest and Inhibits Cell Proliferation of Human Breast Cancer Cells in Vitro through Mitochondria-Mediated Apoptosis. Biol Pharm Bull. 2017;40(8):1232–1239.

18. Ying XJ, Jin B, Chen XW, Xie J, Xu HM, Dong P. Oxymatrine downregulates HPV16E7 expression and inhibits cell proliferation in laryngeal squamous cell carcinoma Hep-2 cells in vitro. Biomed Res Int. 2015;2015:150390.

19. Wang Z, Xu W, Lin Z, Li C, Wang Y, Yang L, et al. Reduced apurinic/apyrimidinic endonuclease activity enhances the antitumor activity of oxymatrine in lung cancer cells. International journal of oncology. 2016;49(6):2331–2340.

20. LiFS, Weng JK. Demystifying traditional herbal medicine with modern approach. Nat Plants. 2017 Jul;3:17109.

